# Perceptual Support for Temporal Normalization Across Hundreds of Milliseconds

**DOI:** 10.1101/2025.08.28.672950

**Authors:** Michael L. Epstein, Sophie Kovacevich, Rachel N. Denison

## Abstract

Perception across time is limited: phenomena like masking and temporal crowding reveal perceptual interference between sequential stimuli, even when separated by hundreds of milliseconds. However the underlying causes of these suppressive temporal interactions remain unclear. A potential explanatory mechanism is temporal normalization, a form of divisive suppression proposed to operate across time in the visual cortex, but largely untested behaviorally. Here in three experiments we tested a key perceptual prediction of temporal normalization: contrast-dependent suppression between sequential stimuli. By independently manipulating the contrasts of two sequential gratings presented 250 ms apart, we found that perceptual sensitivity to a target’s orientation decreased when a high-vs. low-contrast non-target appeared either before or after it. Reduced sensitivity arose from both decreased orientation precision and increased “swapping” errors. The results provide perceptual support for a temporal normalization computation operating across hundreds of milliseconds, offering a theoretical framework for perceptual limits across time.

**Public significance statement:** Temporal context strongly affects perception, but a general theoretical framework that captures such effects is lacking. Neural studies support the existence of temporal normalization, the divisive suppression of information across time. However the behavioral impact of temporal normalization on perception remains unclear. Here we tested a key behavioral prediction of temporal normalization: contrast-dependant suppression between sequential stimuli. We found evidence for such suppression across hundreds of milliseconds both forward and backward in time. Our results provide perceptual evidence for temporal normalization, linking neural data to behavior, and support a theoretical framework that could bridge a range of temporal phenomena such as visual masking, adaptation, and temporal crowding.

## Introduction

Humans dynamically process complex information that continuously unfolds across space and time. However, perception across time is limited, and sequential stimuli can perceptually interfere or otherwise interact with each other across intervals of up to hundreds of milliseconds. For example, in visual masking, a distractor stimulus can disrupt the processing of a target stimulus presented close in time, typically under 150 ms, before or afterwards (Breitmeyer & Öğmen, 2006). In the attentional blink, processing a target stimulus within a rapid sequence can impair identification of a subsequent item (Dux & Marois, 2009). And in adaptation (Webster, 2015) and serial dependence (Fischer & Whitney, 2014), a stimulus can suppress or shift perception of a later one. A more recently described example of perceptual interference across time is temporal crowding (Bonneh et al., 2007; Tkacz-Domb & Yeshurun, 2017, 2021; Yeshurun et al., 2015). In temporal crowding, sensitivity to a target stimulus is reduced when that target is flanked before and after in time by distractors. Notably, temporal crowding occurs for brief sequential stimuli even when the interval between stimulus onsets is as long as 475 ms (Tkacz-Domb & Yeshurun, 2021), outside the range of ordinary masking (Breitmeyer & Öğmen, 2006). To date, no specific mechanistic or computational account of temporal crowding has been proposed. Moreover, although these various temporal phenomena have traditionally been studied within separate literatures, their shared dependence on temporal context—resulting in perceptual interactions between sequential stimuli—raises the question of whether they may be mediated by any common mechanisms.

Here we consider a candidate mechanism that could contribute to a variety of temporal contextual effects in perception: temporal normalization, or divisive normalization that operates across time. Spatial divisive normalization is a well established efficient coding strategy in visual neuroscience, in which the activity of a given neuron is divided by the population activity of neighboring neurons, resulting in a normalized response that efficiently signals changes in visual input across space (Carandini & Heeger, 2012; Heeger, 1992). Evidence of similar normalization computations have been found in multiple neural systems—for sensory coding, spatial attention, reward, and decision making (Louie et al., 2011, 2013; Reynolds & Heeger, 2009)—leading to the proposal that normalization is a “canonical” neural computation (Carandini & Heeger, 2012).

Importantly, normalization has also been proposed to operate across time, and has been linked to temporal interactions between stimuli such as adaptation and masking (Tsai et al., 2012; Webster, 2015). In visual adaptation, for example, normalization provides a succinct explanation for both neural gain and orientation tuning changes (Westrick et al., 2016). Temporal normalization has been particularly successful in explaining cases of temporal suppression of neural responses (Chapman & Denison, 2025). Using MRI, Zhou et al. (2018) found that successive stimuli resulted in a non-linear summation of BOLD responses and could account for these signals using a delayed divisive normalization model. Delayed normalization could also explain electrocorticography data from experiments in which observers viewed stimuli with varying inter-stimulus intervals and contrasts (Brands et al., 2024; Groen et al., 2022; Zhou et al., 2019). Altogether these results support the idea that normalization in the visual system operates across both space and time.

Neural evidence for temporal normalization has been reported at time ranges up to 500 ms (Groen et al., 2022; Zhou et al., 2018), inviting the question of whether and how temporal normalization at these timescales impacts perception. A key property of normalization is the contrast dependence of interactions between stimuli (Carandini & Heeger, 2012). If two stimuli interact via a normalization computation, they will mutually suppress each other, and a higher contrast stimulus will produce more suppression than a lower contrast stimulus. For example, in spatial surround suppression, a high-contrast surround perceptually suppresses the center portion of the stimulus more strongly than a low-contrast surround (Simoncelli & Schwartz, 1999; Zenger-Landolt & Heeger, 2003). Likewise, in adaptation, a high-contrast adapter perceptually suppresses an immediate subsequent test stimulus more strongly than a low-contrast adapter (Greenlee et al., 1991). Here, in three psychophysical experiments, we took advantage of this basic property of normalization computations to test whether temporal normalization is a viable mechanism to explain perceptual interference across hundreds of milliseconds, as in temporal crowding.

We used a simplified temporal crowding protocol in which two Gabor stimuli were presented sequentially in the same location, separated by a 250 ms stimulus-onset asynchrony (SOA). Participants reported the orientation of one of the stimuli, indicated by a response cue, at the end of each trial. Critically we manipulated the contrast of each stimulus independently, allowing us to test whether performance decreased when the irrelevant non-target stimulus was high vs. low contrast, as predicted by temporal normalization. Importantly, given the directionality of time, our two-target protocol also allowed us to test whether and how contrast-dependent temporal suppression occurs forward in time, backward in time, or in both directions. Whereas adaptation is associated with forward suppression (an earlier stimulus suppresses a later one) (Webster, 2015), masking can yield both forward and backward suppression (a later stimulus suppresses an earlier one) (Breitmeyer & Öğmen, 2006), suggesting that across short timescales temporal suppression can be bidirectional. At longer timescales, temporal crowding has always been tested with flankers both before and after the target (Bonneh et al., 2007; Tkacz-Domb & Yeshurun, 2017, 2021; Yeshurun et al., 2015), so its directionality is unknown.

## Experiment 1

### Methods

#### Participants

Eight participants completed Experiment 1 (mean age = 24.8 years, 4 female, all right handed). Two of the participants are authors (ME and SK). The selected sample size was based on the number of participants in previous experiments with similar protocols (i.e., Denison et al., 2017, 2021). All participants had normal or corrected-to-normal vision. All procedures were approved by the Boston University Institutional Review Board.

#### Stimuli

Stimuli were generated on a Dell computer using custom scripts written in MATLAB with Psychtoolbox (Brainard, 1997; Kleiner et al., 2007; Pelli, 1997), and presented on a ViewPixx LCD monitor with a resolution of 1920×1080 pixels. Visual stimuli were Gabor patches constructed as sinusoidal gratings with a spatial frequency of 4 cycles per degree and a 2D Gaussian envelope (0.7° standard deviation) and were displayed in the lower right hand corner of the monitor, 5.7° from a central fixation circle (.45° diameter; see **Figure 1** for an example). Stimuli were presented at either high contrast (64%) or low contrast (4%) and could be tilted either clockwise or counterclockwise about either the vertical or horizontal axis, with independent axes and tilts for each stimulus. The independent axes across stimuli were employed to discourage participants from comparing tilts across sequential stimuli. Auditory stimuli were pure tones, which could be high (784 Hz) or low (523 Hz), or their combination. Tones were enveloped by 10 ms cosine ramps at their start and end, and were presented through speakers at a comfortable volume.

**Figure 1:**
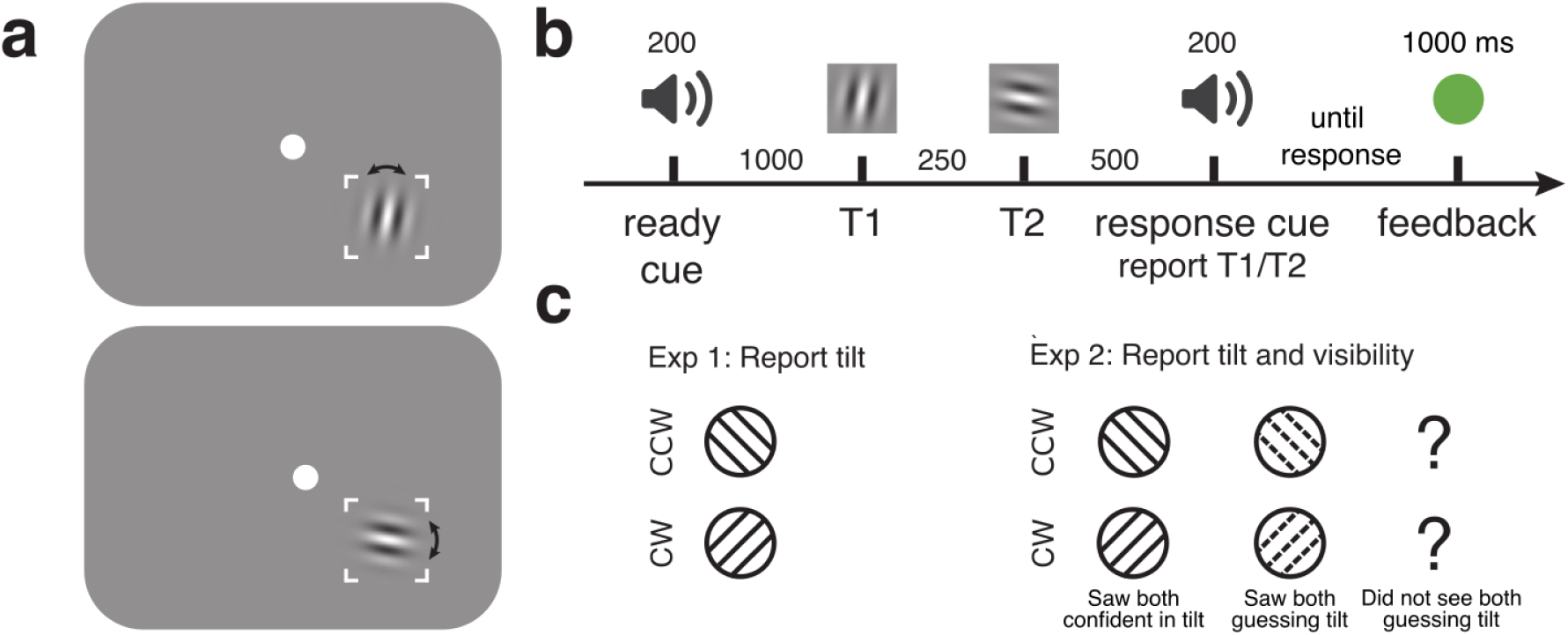
Task design for Experiments 1 and 2. (a) Example stimuli. (b) Task sequence: participants heard a readiness cue, followed 1000 ms later by two stimuli separated by 250 ms. A response cue at the end of the trial instructed participants to report the tilt (clockwise or counterclockwise) of either the first stimulus (T1) or the second stimulus (T2). In Experiment 1, stimulus duration was 100 ms, while in Experiment 2 duration was 50 ms. (c) Possible responses for each experiment. In Experiment 1 only tilt was reported. In Experiment 2 tilt and visibility were simultaneously reported with one key press.

To reduce spatial uncertainty, the screen location where Gabor patches appeared was indicated by lines 0.056° wide showing the corners of a 4.25° x 4.25° square. The fixation color was white, throughout the trial, with color changes used to provide accuracy feedback at the end of each trial: green for a correct response, red for incorrect. Participants sat 57 cm from the screen with their head stabilized in a chin-and-head rest. The testing room was dimly lit.

#### Procedure

On each trial participants viewed two sequentially presented Gabor patches (T1 and T2), each with a 100 ms duration, separated by a 250 ms SOA. A 200 ms response cue tone indicating the target stimulus was presented 500 ms after the onset of T2. A high tone indicated T1 was the target, and a low tone indicated T2 was the target. Participants had unlimited time to report whether the target Gabor was tilted clockwise or counterclockwise from the vertical or horizontal axis. A readiness cue presented 1000 ms before T1 allowed participants to predict the timing of the upcoming stimuli. The readiness cue consisted of simultaneous high and low tones that did not provide any information about which stimulus would be the target. After responding, participants received accuracy feedback via a change in fixation color with a 1000 ms duration. All combinations of stimulus tilts, axes, contrasts, and response-cued targets were presented, randomly intermixed, resulting in 384 trials per session and 1152 trials for the full experiment. One subject only completed 2 sessions, with 768 trials total.

During the experiment, fixation was monitored with an EyeLink 1000 Plus eye tracker (SR Research). Participants were required to maintain fixation for at least 300 ms before a trial could begin. If participants blinked or broke fixation at any time between the readiness and response cues, the trial ended early, and was moved to the end of the trial sequence to be displayed later in the experiment.

#### Training

Experimental sessions were preceded by a training session in which participants learned the task and performed a 5-up 1-down staircasing procedure to find a tilt threshold that achieved approximately 87% accuracy (Leek, 2001). Stimuli in this procedure matched the high contrast (64%) stimuli in the main task. The staircasing procedure used two interleaved staircases to ensure convergence. If either staircase resulted in a tilt threshold of 8° (the maximum tilt included in the staircase), or the staircases didn’t converge, staircasing was repeated. Participants achieved an average 4.75° threshold. At the beginning of each session participants also performed a refresher practice block consisting of 64 trials before starting the main experiment to ensure they were prepared for the task.

#### Analysis

Trials were split into four conditions for each target based on the contrast of the target (high or low) and the contrast of the non-target (high or low). For each of these conditions orientation discrimination performance was calculated as *d’*. All trials were used for this analysis. To calculate statistics, we used 2×2×2 repeated measures ANOVAs with target contrast, non-target contrast, and target (T1 or T2) conditions. Follow-up ANOVAs were run for each target individually, as well as two-tailed paired t-tests for statistics comparing non-target contrast levels for each target and target contrast level. We report confidence intervals on the estimated marginal mean differences modeled by the ANOVAs between conditions showing main effects.

#### Transparency and Openness

Raw data and code to perform our analysis are available at https://osf.io/vf2ac/. This study was not preregistered.

### Results and discussion

If temporal normalization impacts perception, perceptual sensitivity for a target stimulus should be reduced when the target is temporally flanked by a high-vs. low-contrast non-target. We tested this hypothesis by assessing perceptual sensitivity in the orientation discrimination task for each target as a function of both its contrast and the contrast of the non-target, which was irrelevant for the participant’s response (**Figure 2**). Perceptual sensitivity increased with target contrast (estimated marginal mean difference (EMMD) = 1.877, 95% confidence interval = [1.595 2.158], F(1,7) = 248.268, p < 0.001, η_p_^2^ = 0.973), as expected, and was higher for T2 than for T1 (EMMD = 7.46 [0.247 1.246], F(1,7) = 12.479, p = 0.01, η_p_^2^ = 0.641). Importantly, perceptual sensitivity was also impacted by the non-target contrast, with lower orientation discriminability when the non-target contrast was higher (EMMD = 0.766 [0.531 1.001], F(1,7) = 59.580, p < 0.001, η_p_^2^ = 0.895). Additionally, for T1 the impact of the non-target was stronger for high contrast targets, whereas for T2 it was stronger for low contrast targets (3-way interaction between target, target contrast, and non-target contrast, F(1,7) = 7.169, p = 0.032, η_p_^2^ = 0.506).

**Figure 2:**
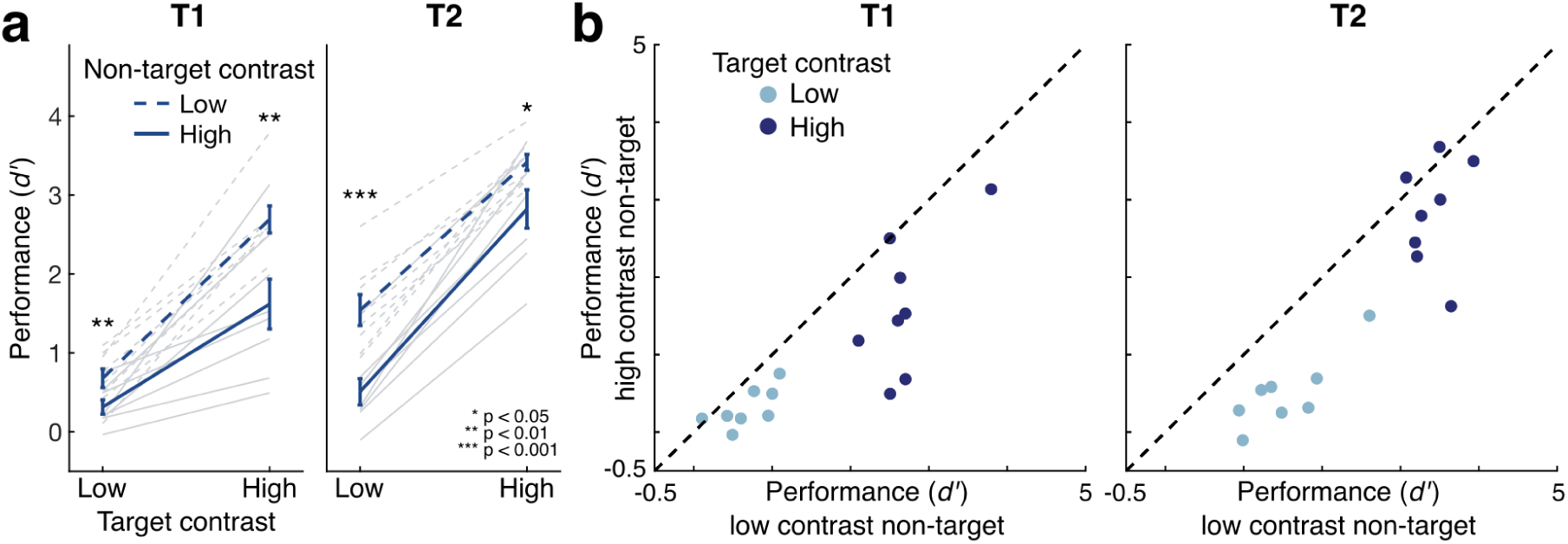
Non-target contrast impacts perceptual sensitivity. (a) Sensitivity to target stimuli increased with higher target contrast, but decreased with higher non-target contrast. Dark lines show group means; light lines show individual participants. Error bars give ±1 SEM. Asterisks show results of pairwise t-tests between high and low non-target contrast conditions. (b) Scatterplots show consistency of effects across individual participants (data points). Points below the unity line exhibit contrast-dependent suppression by the non-target.

We also assessed performance for each target separately. Perceptual sensitivity to T1 increased with target contrast (EMMD = 1.659 [1.513 2.164], F(1,7) = 60.176, p < 0.001, η_p_^2^ = 0.896), and decreased with non-target contrast (EMMD = 0.719 [0.382 1.056], F(1,7) = 25.395, p = 0.001, η_p_^2^ = 0.784). The suppression of perceptual sensitivity by high contrast non-targets was stronger for high-contrast than for low-contrast T1 targets (F(1,7) = 9.625, p = 0.017, η_p_^2^ = 0.579). T2 showed similar results: sensitivity increased with target contrast (EMMD = 2.094 [1.793 2.396], F(1,7) = 269.365, p < 0.001, η_p_^2^ = 0.975) and decreased with non-target contrast (EMMD = 0.813 [0.496 1.130], F(1,7) = 36.788, p < 0.001, η_p_^2^ = 0.840). For T2, the suppression of perceptual sensitivity was similar across target contrasts (F(1,7) = 2.604, p = 0.151, η_p_^2^ = 0.271).

The results show clear contrast-dependent suppression by the non-target at a timescale beyond that typically associated with backwards or forwards masking. Suppression extended both forward and backward in time, with higher-contrast non-targets reducing target discriminability whether the non-target occurred before or after the target. The results are consistent with temporal normalization, which is expected to reduce perceptual sensitivity to a target in the presence of a high-contrast sequential non-target.

## Experiment 2

Experiment 1 provided initial evidence that the kinds of temporal interactions predicted by temporal normalization are indeed present with stimuli separated by 250 ms intervals. However, an alternative possibility is that high-contrast non-targets led participants simply to miss a target, which could have other explanations besides normalization. Perceiving only one of the two stimuli could also create ambiguity when reporting based on the response cue, as participants may be unsure whether the stimulus they saw was T1 or T2. In Experiment 2 we addressed this possibility by increasing the likelihood that participants saw both stimuli during each trial. To do so, we 1) increased the contrast of the lower-contrast stimulus and 2) asked participants to report stimulus visibility along with their tilt judgment, which allowed us to restrict our analysis to trials in which both stimuli were reported as seen.

### Methods

#### Participants

Seventeen participants were recruited for the experiment. Five did not achieve at least 80% accuracy during practice and so did not progress past the practice stage, leaving the targeted number of 12 participants in the final sample (average age 25.4 years, 7 female, 1 left handed, 1 ambidextrous). All participants had normal or corrected-to-normal vision. Five participants for this experiment also participated in Experiment 1, including ME and SK.

#### Stimuli

Stimuli were identical to those in Experiment 1, but with 50 ms visual stimulus durations and contrasts of 64% (high) and 16% (low). The low contrast value was increased 4-fold compared to the value used in Experiment 1 to increase the visibility of the low contrast stimuli, and make it more likely that participants would see both stimuli on each trial.

#### Procedure

In Experiment 2 the procedure was the same as in Experiment 1, except that participants gave a simultaneous visibility judgment along with their tilt report. Visibility reports had three levels, which were designed to prevent participants from conflating confidence about the tilt with the visibility of the grating: 1) “Saw both stimuli and am confident about the target’s tilt”, 2) “Saw both stimuli but am guessing the target’s tilt”, or 3) “Did not see both stimuli (i.e., saw only one or none). Participants were further instructed using example scenarios for each visibility level. For example, a level 3 example was: “I only saw one grating, and so I am unsure which was the target, but I am guessing counter-clockwise.” Participants used a single keypress to report both tilt and visibility. They used the 1, 2 and 3 keys for counterclockwise tilts, and 8, 9 and 0 for clockwise tilts. Outer keys (1,0) were used to report high visibility of both stimuli (level 1), middle keys (2,9) for intermediate visibility (level 2), and inner keys (3,8) for having missed at least one stimulus (level 3).

#### Training

Training was similar to Experiment 1, except that beginning with the contrast training block participants were additionally trained in using the visibility reports. Additionally, thresholding was performed using two interleaved 3-up-1-down staircases to achieve approximately 80% accuracy with high-contrast stimuli. Participants in this experiment had an average 3.58° tilt threshold.

#### Analysis

The analysis was identical to Experiment 1, except that visibility reports were used to filter trials by removing all trials where participants marked at least one stimulus as being missed (visibility level 3; on average 8.73% of trials). This ensured that we were examining only suprathreshold trials in which both stimuli were reported as visible, regardless of the participant’s confidence in their orientation report.

### Results and discussion

Even with the more conservative contrast levels and trial selection procedure employed here, we again found clear suppression of perceptual sensitivity when high-contrast non-targets were present, aligning with predictions of temporal normalization even in suprathreshold cases (**Figure 3**). Perceptual sensitivity increased with higher target contrasts (EMMD = 0.936 [0.737 1.135], F(1,11) = 107.264, p < 0.001, η_p_^2^ = 0.907) and was higher for T2 overall (EMMD = 0.809 [0.453 1.164], F(1,11) = 25.113, p < 0.001, η_p_^2^ = 0.695). Perceptual sensitivity again decreased with higher non-target contrasts (EMMD = 0.388 [0.219 0.558], F(1,11) = 25.415, p < 0.001, η_p_^2^ = 0.698). No interactions reached significance (all F(1,11) < 3.484, p > 0.09).

**Figure 3:**
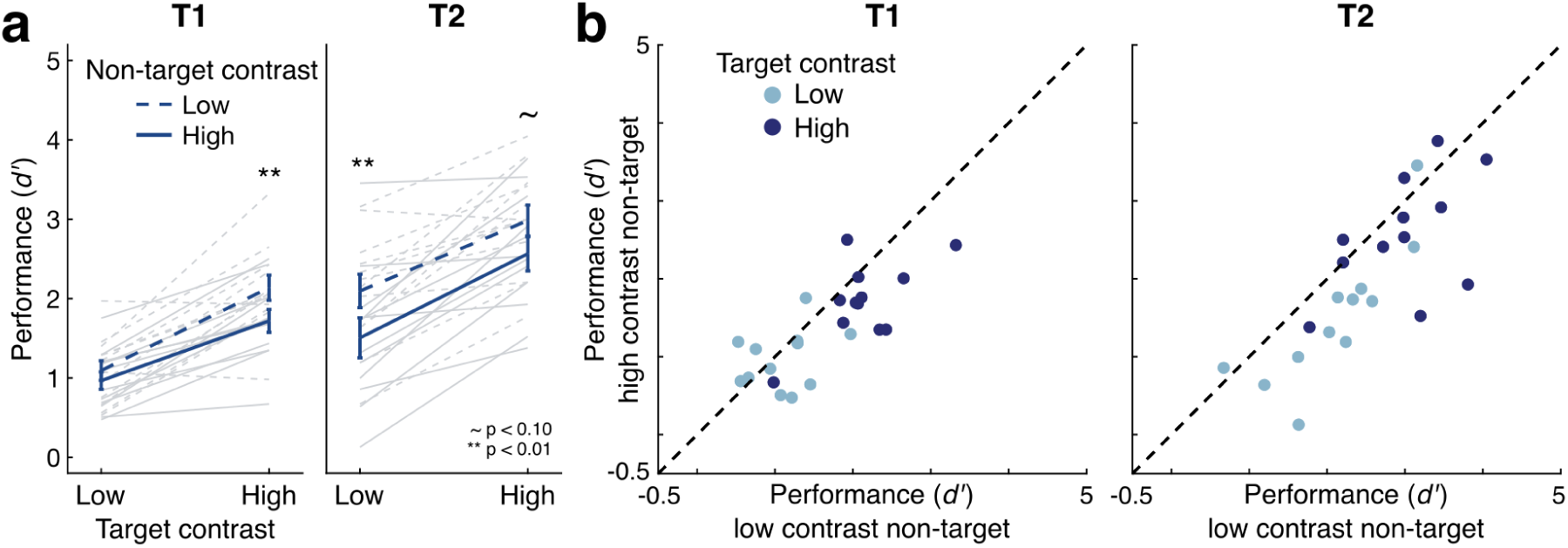
Effects of contrast on perceptual sensitivity in suprathreshold trials. (a) Sensitivity to target stimuli increased with higher target contrast and decreased with higher non-target contrast. Dark lines show group means; light lines show individual participants. Error bars give ±1 SEM. Asterisks show results of pairwise t-tests between high and low non-target contrast conditions. (b) Perceptual sensitivity on trials with low contrast non-targets plotted against sensitivity on trials with high contrast non-targets show that individual participants (data points) consistently fall beneath the unity line, indicating contrast-dependent suppression by non-targets.

We found the same patterns were observed for each target individually. Perceptual sensitivity for T1 increased with higher target contrasts (EMMD = 0.898 [0.672 1.124], F(1,11) = 76.302, p < 0.001, η_p_^2^ = 0.874) and decreased with higher non-target contrasts (EMMD = 0.272 [0.063 0.482], F(1,11) = 8.165, p = 0.016, η_p_^2^ = 0.426). Unlike in Experiment 1, the strength of non-target suppression was consistent across target contrasts (F(1,11) = 2.317, p = 0.156, η_p_^2^ = 0.174), suggesting that the selected contrasts may have better targeted the middle of the dynamic range. Perceptual sensitivity for T2 similarly increased with target contrast (EMMD = 0.973 [0.657 1.290], F(1,11) = 45.790, p < 0.001, η_p_^2^ = 0.806), decreased with non-target contrast (EMMD = 0.504 [0.215 0.794], F(1,11) = 14.704, p = 0.003, η_p_^2^ = 0.572), and showed similar levels of suppression for both high and low contrast targets (F(1,11) = 0.538, p = 0.479, η_p_^2^ = 0.047).

These results further support the existence of contrast-dependent temporal suppression by a non-target both forward and backward in time. This suppression was robust even with the more conservative experimental design, as compared to Experiment 1, with the differences between high and low contrasts reduced 4-fold and only trials reported as visible included in the analysis. These results provide evidence that contrast-dependent temporal suppression occurs across time for suprathreshold stimuli. Together with Experiment 1, it shows that contrast-dependent temporal suppression is strong and consistent across a range of stimulus strengths and task difficulties.

## Experiment 3

The reduction in perceptual sensitivity in the presence of high-contrast non-targets observed in Experiments 1 and 2 could arise in various ways, reflecting different kinds of changes in stimulus processing. It could result from reduced precision in the representation of the target orientation, an increase in random guessing, reporting the non-target instead of the target (swapping), biasing one’s responses towards the orientation of the non-target, or a combination of these factors.

Temporal normalization makes the clearest prediction regarding precision: if normalization occurs across time, high contrast non-targets should reduce the strength of target representations, resulting in less precise orientation estimates even when the target remains visible. An observed loss of precision would provide evidence in support of normalization while controlling for other potential effects of the temporal interaction. At the same time, determining the contribution of multiple potential causes of contrast-dependent temporal suppression in performance accuracy offers the opportunity to better understand this phenomenon.

To more fully understand how temporal suppression affects the stimulus information available to the participant, we conducted a third experiment which was identical to Experiment 2, except we replaced the two-choice orientation discrimination task with a continuous orientation estimation task. Stimuli were presented with random orientations, and we asked participants to report the orientation of the target stimulus using an adjustable response probe. By fitting probabilistic mixture models to the distributions of errors, we tested the extent to which high-contrast non-targets led to a decrease in the quality of the orientation representations (resulting in a loss of precision), a more complete loss of target orientation information (resulting in increased guess rates or reports of the non-target orientation), or temporal integration between the target and non-target representations (resulting in bias to the non-target). Thus the orientation estimation task allows a more detailed picture of how the observed temporal suppression influences the available stimulus information.

### Methods

#### Participants

Twenty-six participants were recruited for Experiment 3. Six did not achieve better than an average 20° error during training, and so did not progress to the main experiment. This left the targeted number of 20 participants in the final sample (average age 25.1 years, 15 female, 1 left handed, 1 ambidextrous). All participants had normal or corrected-to-normal vision.

#### Stimuli

Stimuli were identical to those in Experiment 2, except that the stimulus orientations of T1 and T2 were now independently and randomly selected from 0-179° in steps of 1°. The orientation response probe was a single white line (3° in length, 0.1° width) presented in the same location as the grating stimulus. The probe could be rotated by moving the mouse left and right.

#### Procedure

The procedure was similar to Experiment 2, but adapted for the orientation estimation task. Participants were shown an orientation response probe simultaneously with the response cue, in the same location where the stimuli were presented. Participants could adjust the orientation of the response probe by moving the mouse left and right and were instructed to match the orientation of the probe as closely as possible to the orientation of the target stimulus before clicking the mouse to submit their response. They were given unlimited time to respond. Participants used a left or right click to report one of two visibility levels on each trial: a left click indicated that both targets were visible (level 1), and a right click indicated that one or both targets were missed (level 2). Participants received accuracy feedback after each trial, with their error (rounded to the degree) presented for one second on the screen 1° above fixation. On trials where the participant’s error was low, they additionally received positive feedback (<10°: “Good!”, <5°: “Great!”, <1°: “Nailed it!”). The response-cued target and stimulus contrasts were balanced and randomly intermixed across trials. We collected 640 trials over two sessions for each participant.

#### Training

The training procedure was the same as for Experiment 2, but with the criterion for advancing to the main experiment modified for the estimation task. Participants had to achieve less than 20° average error during practice to proceed to the main experiment.

#### Analysis

On each trial we calculated a participant’s orientation estimation error as the angle between the reported orientation and the target orientation. Similarly as in Experiment 2, we removed trials where participants had marked at least one stimulus as “missed” (visibility level 2; on average 14.91% of trials). To test the effects of the temporal context on orientation estimation, we employed a probabilistic mixture-model, drawn from recent work exploring working memory and serial dependence (Moon, 2025; Moon et al., 2024). This model offers some advantages over classic mixture models (i.e., Bays et al., 2009; Gorgoraptis et al., 2011; Zhang & Luck, 2008) in that in addition to fitting distributional parameters (such as the standard deviation of responses to the targets) it further calculates the probability that each trial reflected a response to the target, the non-target, or a random guess. This approach can improve estimates of how much orientation responses are biased to a non-target by restricting the analysis to trials in which the participant was most likely responding to the target. Without this step, estimates of bias are generally inflated by mixing in trials in which participants were likely responding to the non-target (Moon, 2025; Moon et al., 2024). This model was well-suited to our experimental design, as we had just two stimuli per trial, which could plausibly give rise to either swapping or integration.

To implement this model we fit von Mises distributions to error relative to the target and non-target orientations along with a uniform guess rate parameter. The probability of a given orientation response *R* is modeled as

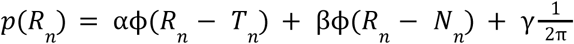

where *T* is the target stimulus orientation and *N* is the non-target orientation on trial *n*. Responses to the target and non-target are modeled as being drawn from von Mises distributions ϕ centered at zero with a fitted standard deviation, σ, of the circular distribution, and evaluated at the value given in parentheses in the terms of the equation. α is the probability that a given response was based on the target, β that it was based on the non-target, and γ that it was a random guess drawn from a uniform distribution across orientation. Fitted parameters were thus: α, β and σ. The parameter γ was calculated as 1 − α − β. We fitted this model to trial data for each condition and participant separately.

Over 1000 iterations we fit this model to the error patterns in the data. On each iteration we then compared the error for each trial to the von Mises probability density functions of each of these fits, to calculate the likelihood that the error could be explained by each response type. By normalizing the sum of these probabilities, along with the likelihood of pure guessing, to 1, we can construct a ratio of the probabilities of each response type for that specific trial. Then by generating a random number from the uniform distribution between 0 and 1, we can tag the trial by which response type it fell into. Over the 1000 iterations, the average number of times a given trial is sorted into each category can be used to calculate the probability that that trial belonged to that category.

With these results, we could extract both the standard deviation (SD) parameter (which provides the inverse of target precision) as well as the percentage of trials labelled by the model as swaps or guesses, averaged across the 1000 iterations of model fitting.

We used repeated measures ANOVAs to calculate separate statistics for precision, swap rate, and guess rate. Finally, by selecting trials where target responses were most likely, we estimated bias to the non-target by calculating the error relative to the target plotted against the error relative to the non-target, again averaged across 1000 iterations of model fitting. These values were compared to zero to test overall bias and compared across conditions to test the effects of target and non-target contrast on bias, both using cluster-corrected permutation tests to control for type-1 errors (Maris & Oostenveld, 2007).

### Results and discussion

Target performance decreased in the presence of high-vs. low-contrast non-targets, confirming the findings of Experiments 1 and 2 with a different type of orientation report. Orientation estimation performance followed a pattern similar to discrimination performance in the other experiments: error was greater when non-targets were higher contrast (EMMD = 2.660° [1.539 3.780], F(1,19) = 24.689, p < 0.001, η_p_^2^ = 0.565), decreased as target contrast increased (EMMD = 4.136° [2.507 5.766], F(1,19) = 28.224, p < 0.001, η_p_^2^ = 0.598), and was lower overall for T2 compared to T1 (EMMD = 7.088° [4.548 9.629], F(1,19) = 34.100, p < 0.001, η_p_^2^ = 0.642), (**Figure 4a**). Relative to T2 performance, T1 performance improved more as target contrast increased (interaction between target contrast and target: F(1,19) = 28.968, p < 0.001, η_p_^2^ = 0.604) and exhibited stronger suppression by non-target contrasts (interaction between non-target contrast and target: F(1,19) = 6.931, p = 0.016, η_p_^2^ = 0.267). Examining these effects separately for each target showed that for T1, error decreased with target contrast (EMMD = 5.762° [3.773 7.752], F(1,19) = 36.751, p < 0.001, η_p_^2^ = 0.959) and increased with non-target contrast (EMMD = 3.273° [1.947 4.598], F(1,19) = 26.698, p < 0.001, η_p_^2^ = 0.584). The strength of non-target suppression was similar across target contrasts (F(1,19) = 2.215, p = 0.153, η_p_^2^ = 0.104). For T2, error also decreased with target contrast (EMMD = 2.510° [1.043 3.977], F(1,19) = 12.8222, p = 0.002, η_p_^2^ = 0.403), and increased with non-target contrast (EMMD = 2.047° [0.683 3.410], F(1,19) = 9.865, p = 0.005, η_p_^2^ = 0.342). A trending interaction suggests that for T2 the effect of the non-target was stronger for low-contrast compared to high-contrast targets (F(1,19) = 4.236, p = 0.054, η_p_^2^ = 0.182).

**Figure 4:**
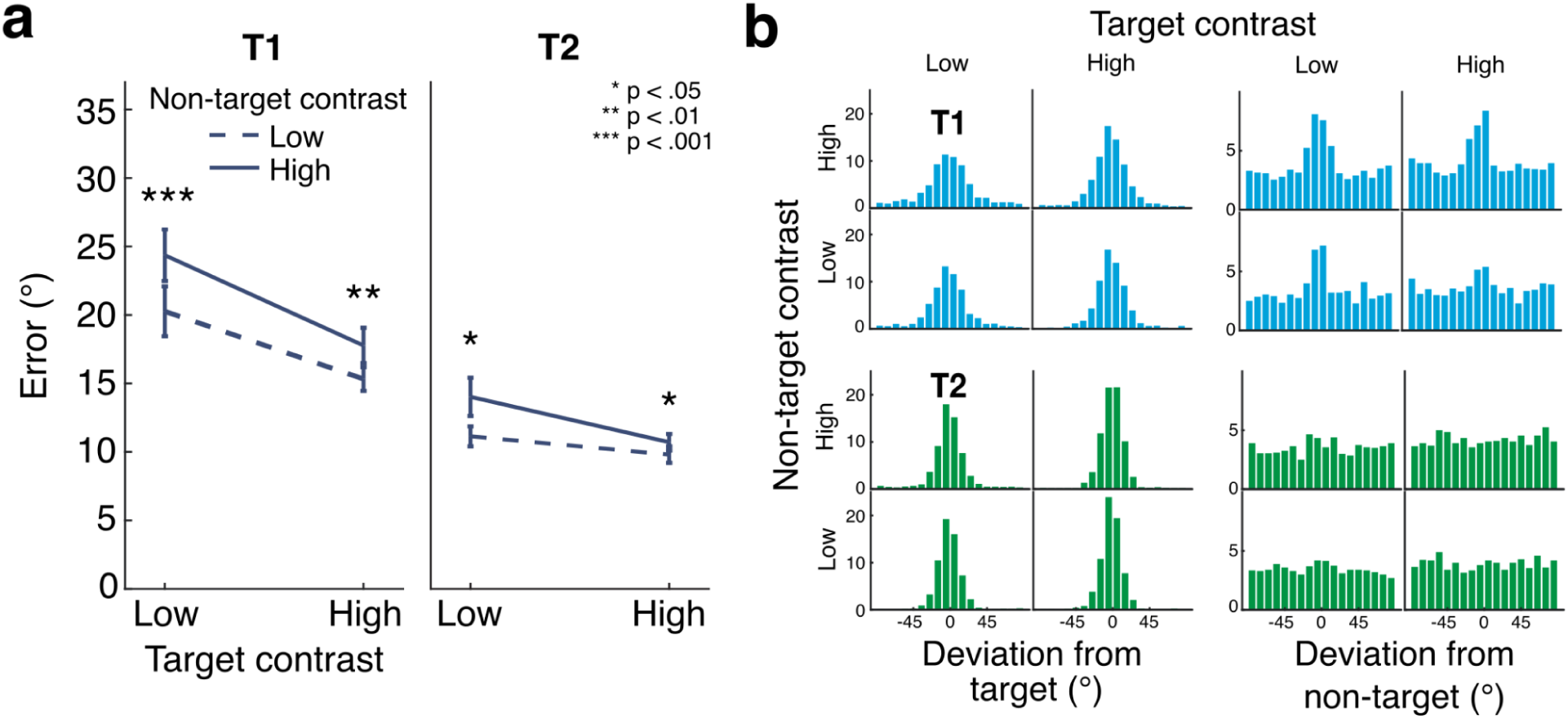
Orientation estimation task. (a) Average error across participants shows a similar pattern to that seen in Experiments 1 and 2: error increased with non-target contrast and decreased with target contrast. Error bars give ±1 SEM. Asterisks show results of pairwise t-tests between high and low non-target contrast conditions. (b) Histograms (mean trial counts across participants) showing distributions of response deviations from target (left) and non-target (right) orientations.

Next we visualized the distribution of errors, both with respect to the target and the non-target stimulus orientations. Most orientation reports were distributed around the target orientation, as expected, but small peaks in the error distributions relative to the non-target indicate some evidence for swapping as well (**Figure 4b**).

Probabilistic modeling of trial-by-trial orientation reports revealed various contributions to T1 and T2 errors. As an overview, higher contrast non-targets predominantly reduced precision (increased standard deviation, SD) for T1, with some evidence of increased swapping, whereas for T2, higher contrast non-targets predominantly increased swapping (**Figure 5**). After accounting for swaps, there was little additional random guessing for either target. Finally, orientation estimates were biased in the direction of the non-target, but this bias did not depend on the contrast of the non-target.

**Figure 5:**
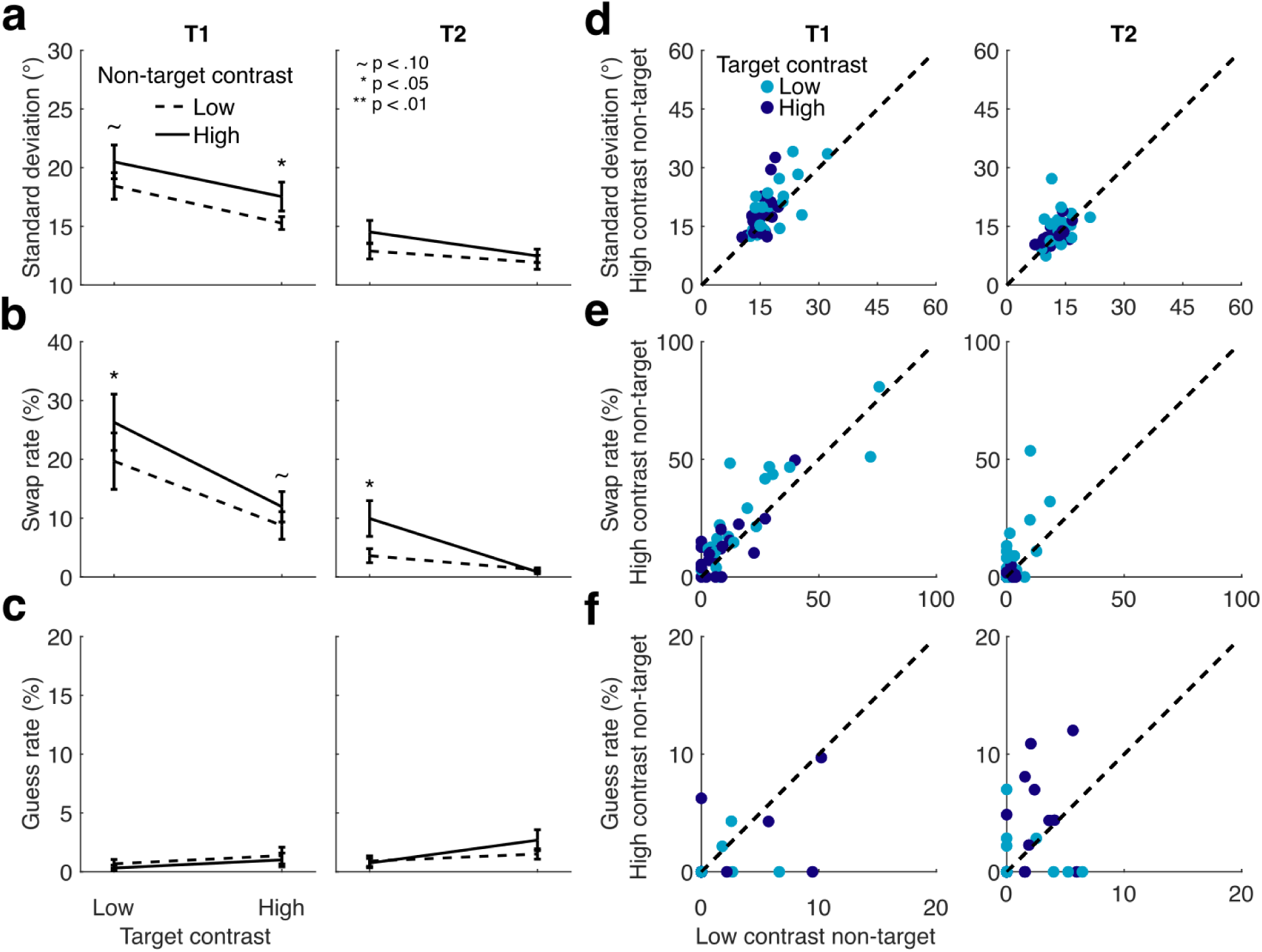
Probabilistic modeling fits to estimation data. (a) Standard deviation increases with non-target contrast for T1. (b) Swap rate increases for both targets at low target contrasts. (c) Guess rate shows no significant effects of non-target contrast. Asterisks show results of pairwise t-tests between high and low non-target contrast conditions. (d-f) Scatterplots show effects across individual participants (data points).

First, the quality of orientation information for each target was reflected in the precision of the orientation estimates for that target (**Figure 4a,d**). In the full ANOVA, SD decreased with higher target contrasts (EMMD = 2.28° [1.214 3.346], F(1,19) = 20.031, p < 0.001, η_p_^2^ = 0.513) and increased with higher non-target contrasts (EMMD = 1.618° [0.479 2.758], F(1,19) = 8.835, p = 0.008, η_p_^2^ = 0.317) across targets. SD was lower for T2 than for T1, similar to the pattern seen for sensitivity in Experiments 1 and 2 (EMMD = 4.976° [3.416 6.537], F(1,19) = 44.554, p < 0.001, η_p_^2^ = 0.701). An interaction revealed the effect of target contrast to be stronger for T1 than for T2 (F(1,19) = 6.854, p = 0.017, η_p_^2^ = 0.265). No other interactions were observed (all F(1,19) < 1.1, p > 0.293, η_p_^2^ < 0.058).

Restricting our tests by target showed that for T1, SD decreased with target contrast (EMMD = 3.060° [1.516 4.605], F(1,19) = 17.205, p < 0.001, η_p_^2^ = 0.475) and increased with non-target contrast (EMMD = 2.154° [0.322 3.986], F(1,19) = 6.053, p = 0.024, η_p_^2^ = 0.242). The effects of non-target contrast on SD were similar across target contrasts (F(1,19) = 0.046, p = 0.832, η_p_^2^ = 0.002). For T2, SD also decreased as target contrast increased (EMMD = 1.5° [0.683 2.317], F(1,19) = 14.760, p = 0.001, η_p_^2^ = 0.437). T2 showed a trending effect of non-target contrast on SD (EMMD = 1.083° [-0.096 2.261], F(1,19) = 3.697, p = 0.07, η_p_^2^ = 0.163. There was no interaction between target and non-target contrast for T2 (F(1,19) = 0.955, p = 0.241, η_p_^2^ = 0.048).

Second, orientation reports could reflect swapping, or reporting the orientation of the non-target (**Figure 4b,e**). Swap rates were higher overall when T1 was the target (EMMD = 12.763% [6.167 19.360], F(1,19) = 16.401, p < 0.001, η_p_^2^ = 0.463), and they decreased with higher target contrast (EMMD = 9.193% [4.720 13.667], F(1,19) = 18.500, p < 0.001, η_p_^2^ = 0.493), more so for T1 than for T2 (F(1,19) = 10.255, p = 0.005, η_p_^2^ = 0.351). Non-target contrast affected swap rates, with higher contrasts increasing swap rates overall (EMMD = 3.934 [2.145 5.723], F(1,19) = 21.174, p < 0.001, η_p_^2^ = 0.527). However, swap rates increased to a greater degree for low-contrast targets as compared to high-contrast targets (interaction between target contrast and non-target contrast, F(1,19) = 10.634, p = 0.004, η_p_^2^ = 0.359). Other effects were not significant (F < 0.804, p > 0.381).

When restricting the ANOVA to T1, higher target contrast reduced swap rates (EMMD = 12.647% [6.731 18.564], F(1,19) = 20.018, p < 0.001, η_p_^2^ = 0.513), and higher non-target contrast increased them (EMMD = 4.891% [1.744 8.038], F(1,19) = 10.582, p = 0.004, η_p_^2^ = 0.358). The effect of non-target contrast did not differ significantly across target contrasts for T1 (F(1,19) = 1.391, p = 0.253, η_p_^2^ = 0.068). Similarly for T2, higher target contrast reduced swap rates (EMMD = 5.739% [1.839 9.640], F(1,19) = 9.484, p = 0.006, η_p_^2^ = 0.333), and higher non-target contrasts increased them (EMMD = 2.977% [0.430 5.524], F(1,19) = 5.983, p = 0.024, η_p_^2^ = 0.239). However, swap rates fell close to zero when T2 was high contrast, eliminating the effect of non-target contrast and resulting in a significant interaction between target contrast and non-target contrast (F(1,19) = 6.821, p = 0.017, η_p_^2^ = 0.264).

Third, we assessed guess rate, reflecting random orientation estimates categorized as unrelated to either the target or non-target orientations (**Figure 4c,f**). The probabilistic model assigned very few trials (1.16% overall) to the guess distribution. Removing trials in which participants reported one or more stimuli as not seen could have contributed to this low value. Even so, a trending effect of guess rate increasing with target contrast was observed (EMMD = 0.984% [-0.024 1.992], F(1,19) = 4.179, p = 0.055, η_p_^2^ = 0.180), as well as a weak trending effect of a higher guess rate for T2 as compared to T1 (EMMD = 0.617% [-0.122 1.356], F(1,19) = 3.053, p = .097, η_p_^2^ = 0.138). All other effects were non-significant (all F < 1.677, p > 0.211). When follow-up tests were restricted by target, guess rate was similar across T1 target and non-target contrast conditions (all F < 1.178, p > 0.291). For T2, guess rate increased slightly but significantly with target contrast (EMMD = 1.276% [0.223 2.329], F(1,19) = 6.428, p = 0.02, η_p_^2^ = 0.253) but was unaffected by the non-target contrast, and showed no interaction between target and non-target contrast (both F < 1.696, p > 0.208). Thus non-target contrast had no significant impact on guess rate.

Finally, we assessed the degree to which orientation reports were biased in the direction of the non-target, specifically for trials in which responses were most likely to the target stimulus (**Figure 6**). Responses were biased in the direction of the non-target for all conditions for T1 (p < 0.05, cluster corrected) but not in any condition for T2 (p < 0.05, cluster corrected). The degree of bias did not significantly differ as a function of non-target contrast (no clusters with p < 0.05), suggesting that performance impairments with high-contrast non-targets did not arise from increased integration with the non-target.

**Figure 6:**
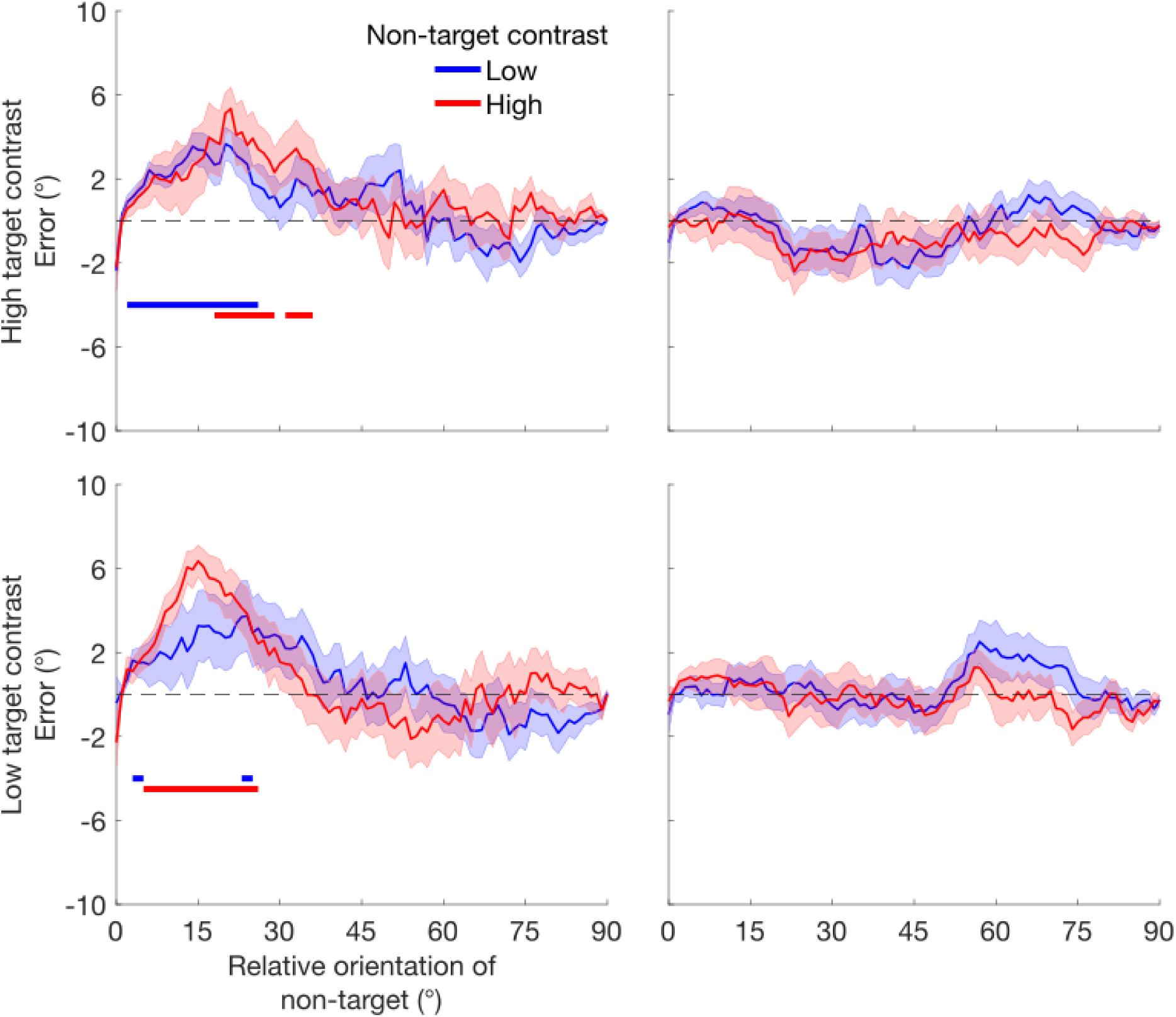
Error biased in the direction of the non-target, restricted to trials where participants were responding to the target according to the probabilistic model. Signed response error (group mean with SEM shown in error ribbon) is plotted against the absolute difference between target and non-target orientations. The horizontal lines mark clusters where bias was significantly different from zero (p < 0.05, cluster-corrected). Responses to T1 show an overall attractive bias to T2, whereas responses to T2 do not show an attractive bias to T1.There were no effects of target or non-target contrast on the levels of measured bias.

Altogether the results of Experiment 3 confirm the presence of contrast-dependent temporal suppression across hundreds of milliseconds and suggest that this suppression arises from a combination of causes. Suppression backwards in time—from T2 to T1—reduced the precision of orientation estimates, consistent with predictions of temporal normalization. Suppression in both directions increased the rate of swap errors, particularly at low target contrasts. Bias to the non-target on T1 trials and low rates of guessing were also observed in our data, but neither was significantly affected by the non-target contrast. Therefore contrast-dependent temporal suppression arises from a combination of reduction in orientation precision regardless of target contrast and an increased likelihood to report the non-target orientation specifically when the target contrast is low.

### General discussion

Here we tested the hypothesis that temporal normalization shapes perception by contributing to contextual interactions across time. Just as spatial normalization predicts contrast-dependent interactions across space (Carandini & Heeger, 2012; Simoncelli & Schwartz, 1999), temporal normalization predicts that the contrast of a non-target will influence perception of a target even when the two stimuli are presented sequentially. The results of three psychophysical experiments supported this prediction. Experiment 1 showed that a higher-contrast non-target reduces perceptual sensitivity to a sequential target. Our design allowed us to separately investigate the predictions of normalization both forward and backward in time, which revealed contrast-dependent suppression in both directions. Experiment 2 confirmed these results even for trials in which participants reported both stimuli as visible, showing that contrast-dependent interactions occur for suprathreshold stimuli. Experiment 3 showed that temporal suppression reduces perceptual precision even for high-contrast stimuli, at least in the backward direction. Together these results are consistent with predictions from temporal normalization extending across hundreds of milliseconds, supporting normalization as a possible mechanism influencing perceptual interactions across time.

Experiment 3, in which we used an estimation task and probabilistic modeling to tease apart the underlying causes of the observed temporal suppression, revealed that suppression was associated with both reductions in orientation precision and increases in reporting the non-target, or swapping. Whereas T1 precision was significantly reduced by high-contrast T2 non-targets, the effect of T1 on T2 precision was only trending, perhaps indicating an asymmetry between backward versus forward suppression. Alternatively, given that precision was higher for T2 overall, it is possible that forward suppression was masked by ceiling performance in the current study. Both targets exhibited increased swapping when they were presented at low contrast in the presence of high-contrast non-targets, despite participants marking both stimuli as visible. Temporal normalization straightforwardly predicts that contrast-dependent suppression would reduce the precision with which a stimulus is represented, and could predict increases in swapping if feature information is strongly suppressed on some trials, leading participants to report the more discriminable high contrast non-target. Another possible account of increased swapping, unrelated to normalization, is that the higher contrast stimulus may drive the decision on some trials, particularly if participants have poor temporal order information.

Altogether, these data demonstrate contrast-dependent suppression across a 250 ms time interval, both forward and backward in time. If temporal normalization mediates these effects, it too would have to operate both forward and backward in time and across this timescale. At a theoretical level, bidirectional temporal normalization would be useful for incorporating temporal context to efficiently code time-varying input. However, most work on temporal normalization has only considered the forward direction—in which previous stimuli suppress subsequent ones—as in adaptation (Webster, 2015; Westrick et al., 2016). For example, dynamic normalization models propose that neural responses are normalized by their own stimulus history, leading to forward suppression, a consequence of the forward directionality of time itself. Recent theoretical work, however, has shown that combining spatial and temporal normalization can generate bidirectional temporal suppression (Chapman & Denison, 2025), consistent with the current data. In this dynamic spatiotemporal attention and normalization model (D-STAN), forward suppression arises from normalization by past responses, and backward suppression arises when ongoing responses to stimuli are suppressed by overlapping responses to later stimuli. Suppression of these later portions of the ongoing responses would be expected to reduce precision, as we observed for T1. The durations of forward and backward suppression in this model depend on the time constants of spatiotemporal receptive fields.

While the observed contrast-dependent suppression supports temporal normalization in perception, temporal processing is complex with multiple concurrent processes. For example, recency and primacy effects differentially affect the quality of how items are represented based on where they are in a sequence and what feature is being reported (Hubert-Wallander & Boynton, 2015). Here we controlled for these influences by always presenting two targets and always comparing performance for a given target to itself under different conditions. Thus the effect of temporal context could be assessed for each target, although other aspects of T1 and T2 processing may differ. Attention may also have varied with the sequential position or the contrast of the stimuli, affecting responses. The role of attention could be examined in future work by manipulating it experimentally. Another process that may contribute to temporal contextual effects is ensemble averaging of sequential stimuli (Albrecht & Scholl, 2010). We observed bias toward the non-target for T1, suggesting some degree of averaging; however, we found no evidence that this bias depended on the non-target contrast. This finding supports the interpretation that temporal normalization is a distinct phenomenon from temporal averaging. Still, it remains an interesting open question how the mechanisms underlying ensemble perception and normalization may be linked, as both are hypothesized to depend on pooled neural activity (Robinson & Brady, 2023; Utochkin et al., 2023).

The bidirectional temporal suppression we observed extended over hundreds of milliseconds, exceeding the range typical of masking (Breitmeyer & Öğmen, 2006) but consistent with that of temporal crowding—the decreased sensitivity to targets when they are temporally flanked by irrelevant distractors up to half a second away (Bonneh et al., 2007; Tkacz-Domb & Yeshurun, 2017, 2021; Yeshurun et al., 2015). Our observation of bidirectional contrast-dependent temporal suppression at this timescale shows that temporal crowding likely arises from both forward and backward suppressive effects and raises the possibility that temporal normalization could underlie temporal crowding. Indeed, the current results suggest that a temporal normalization computation could contribute to a variety of temporal contextual effects in perception (Tsai et al., 2012; Westrick et al., 2016), and future research should determine whether and how it does so. An important step toward this goal will be to determine the length of the window across which contrast-dependent suppression extends in each direction in time, as well as further testing of directional asymmetries. Our task provides a protocol that can be adapted to answer such questions as well as extended to neuroimaging contexts to further develop an understanding of temporal normalization (Brands et al., 2024; Groen et al., 2022; Zhou et al., 2018, 2019).

Altogether, our results show contrast-dependent temporal suppression both forward and backward in time at timescales outside the range of ordinary visual masking. These findings provide initial evidence that temporal normalization impacts perception. This work thus ties together theoretical and neuroscience studies on temporal normalization with well-established temporal perceptual effects that have generally not been discussed within a normalization framework. The perceptual link demonstrated here supports normalization as a critical computation for sensory processing in both space and time. Further work is required to fully map the dynamics of temporal normalization and determine whether and how it may tie together disparate computational and cognitive theories of the temporal dynamics of perception. Initially though, these findings contribute to understanding how the brain efficiently processes dynamic information in a continuously changing world.

### Constraints on Generality

Our study population was primarily students at Boston University. Because we are testing contrast-dependent suppression, a low-level visual process, we would expect our results to hold across the general population. However, it is possible that there could be differences in the temporal interactions we observed based on culture, age or other factors that could influence temporal sensitivity or contextual processing. We have no reason to believe that such other factors would impact our results.

